# Second messenger c-di-AMP regulates multiple antibiotic sensitivity pathways in *Mycobacterium smegmatis* by discrete mechanisms

**DOI:** 10.1101/2023.07.05.547667

**Authors:** Aditya Kumar Pal, Dipankar Ghorai, Xueliang Ge, Biplab Sarkar, Amit Kumar Sahu, Vikas Chaudhary, Suparna Sanyal, Mahavir Singh, Anirban Ghosh

## Abstract

One of the debilitating causes of high mortality in the case of tuberculosis and other bacterial infections is the resistance development against standard drugs. There are limited studies so far to describe how a bacterial second messenger molecule can directly participate in distinctive antibiotic tolerance characteristics of a cell in a mechanism-dependent manner. Here we show that intracellular cyclic di-AMP (c-di-AMP) concentration can modulate drug sensitivity of *Mycobacterium smegmatis* by directly interacting with either a protein effector or with the 5’-UTR regions in mRNA of the genes and thus causing transcriptional downregulation of important genes in the pathways. We studied four antibiotics with different mechanisms of action: rifampicin, ciprofloxacin, erythromycin, and tobramycin and subsequently found that the level of drug sensitivity of the bacteria is directly proportional to the c-di-AMP concentration inside the cell. Further, we unraveled the underlying molecular mechanisms to delineate the specific genes and pathways regulated by c-di-AMP and hence result in differential drug sensitivity in *M. smegmatis*. Thus, our findings of c-di-AMP messenger controlling drug resistance phenotypes of mycobacteria against four different classes of antibiotics is a unique observation that will contribute to scientific advancement in the field.

**One Sentence Summary:** Here we describe how second messenger c-di-AMP modulates the antibiotic sensitivity and resistance profile of *M. smegmatis* by diverse mechanisms.

## Main Text

Antimicrobial resistance poses a huge threat to human civilization where bacterial infections cannot be cured using the existing antibiotics arsenal. Over the years, significant efforts have been made to find out the specific cellular factors and pathways responsible for antibiotic resistance in bacteria (*1*–*3*). Previously it has been found that second messenger (p)ppGpp could induce antibiotic resistance and persistence in many bacteria by diverse mechanisms (*4*–*9*). However, knowledge and understanding about how other second messenger molecules could impact the antibiotic sensitivity and resistance phenotypes of a cell in a target-dependent manner is still lacking. We are interested in finding novel cyclic di-AMP (c-di-AMP) driven drug phenotypes in mycobacteria (*10*) and we have probed the mechanistic role of second messenger c-di-AMP determining drug resistance phenomenon in *Mycobacterium smegmatis* (*11*), which is often used as a surrogate to *Mycobacterium tuberculosis* to evaluate drug leads (*12, 13*). Though there were some previous studies of c-di-AMP regulating beta-lactam resistance in Gram-positive bacteria (*14*–*17*), the mechanism was not clear, and the effect seems to be indirect. Throughout this study, we used two previously characterized *M. smegmatis* strains from our laboratory with altered c-di-AMP levels: c-di-AMP null mutant (Δ*disA*) and a c-di-AMP over-expression mutant (Δ*pde*) along with the WT and respective complemented strains. A previous study (*10*) described how the overproduction of c-di-AMP makes cells sensitive to rifampicin and vancomycin. In this study, we seek to understand the specific underlying mechanisms linking high c-di-AMP concentration with antibiotic sensitivity with rifampicin. In addition to that, we asked if the absence/lack of c-di-AMP also makes cells sensitive/ resistant to any antibiotics and we found that Δ*disA* mutant becomes sensitive to erythromycin, tobramycin and moderately resistant to ciprofloxacin by diverse mechanisms. Altogether, our study highlighted three different c-di-AMP regulated pathways in *M. smegmatis* leading to differential drug sensitivity of the strain.

### c-di-AMP interacts with *M. smegmatis* Arr protein and prevents ribosylation of rifampicin

As mentioned earlier *(10)*, we reported *M. smegmatis* Δ*pde* strain shows higher sensitivity to rifampicin (inhibits bacterial RNA polymerase) compared to *M. smegmatis* WT by 8 fold (MIC: 2 μg/ml.) and in Δ*pde+*pPde strain, the MIC value reverts to WT level (16 μg/ml.). Conversely, we also observed that the overexpression of DisA (pMV261-*disA*) protein (c-di-AMP synthetase) results in an increase in rifampicin sensitivity (4 fold) in *M. smegmatis* confirming the direct role of high c-di-AMP concentration behind the phenotype (**Table 1**). Since DisA is a dual-functional protein in several bacteria (*18, 19*), next, we wanted to confirm that it is c-di-AMP and not the DisA enzyme itself that is responsible for rifampicin sensitivity. To prove that, we overexpressed the catalytically inactive mutant of DisA (D84A) in *M.smegmatis* Δ*disA* background and found no change in MIC compared to empty vector control (16 μg/ml.) (**Table 1**). As Arr was known to be involved in rifampicin ribosylation and acquiring partial resistance against rifampicin (*20*–*22*), we hypothesized that c-di-AMP (at high concentration) could interfere with Arr (Msm_Arr_) enzyme’s function and thus make cells sensitive to rifampicin. To explore that, first, we overexpressed Arr protein (pMV261-*arr)* in Δ*pde* strain we found a 16-fold increase in the MIC value (from 2 μg/ml. to 32 μg/ml.), whereas no such modulation in MIC value was observed with Arr overexpression in WT background (**Table 1**). To prove that the Arr is the key downstream effector molecule, we tested the rifampicin sensitivity of the *M.smegmatis* Δ*arr* +pMV261-*disA* strain and we found, unlike the WT background no difference in MIC was observed compared to blank vector control (**Table 1**). Both these observations directly implied the involvement of c-di-AMP with Arr and thus mediating the rifampicin sensitivity of the cells. Next, we asked whether it was Arr’s expression or catalytic activity that was inhibited by c-di-AMP; and the fact that our RNA-seq data did not point out any significant differencein *arr* expression at the mRNA level in Δ*pde* strain (log2 FoldChange is -0.082), we focused on the Arr activity part. Structural comparison of c-di-AMP and NAD^+^ (Arr enzyme’s substrate) revealed some degree of similarity (both the molecules share an adenine and two ribose sugar moieties in their structure) and we wanted to explore the possible binding of the c-di-AMP molecule to Arr enzyme’s catalytic domain and thus inhibiting the ribosylation activity. First, Arr was overexpressed in *E. coli* BL21 Rosetta (DE3) cells and was purified using Ni-NTA affinity column chromatography followed by size exclusion chromatography (SEC). The SEC profile showed that the protein elutes as a monomer with a molecular weight of ∼16 kDa (**Fig. S1A**). The eluted fractions were analyzed for purity by SDS-PAGE (**Fig. S1B**). Purified protein was further characterized using CD and NMR spectroscopy. CD spectrum of Arr showed that it is mainly α-helical in solution (**Fig. S1C**). This was confirmed by recording a 2D ^1^H-^15^N HSQC NMR spectrum of uniformly ^15^N labelled Arr which showed a spectrum with well-dispersed resonance cross peaks indicating that the protein is well-ordered in solution (**Fig. S1D**). The interaction of Arr with NAD+ and c-di-AMP was monitored using NMR spectroscopy. For the NMR titrations, the unlabelled compounds were titrated into a solution of uniformly ^15^N labelled Arr and a 2D ^1^H-^15^N HSQC NMR spectrum was recorded at each step of titration (**Fig. 1A & 1B**). In the case of both NAD+, c-di-AMP titrations, at a 1:2 protein-to-compound ratio, we observed resonance peak disappearances and/or chemical shift perturbations of several peaks in the 2D ^1^H-^15^N HSQC NMR spectrum of Arr (**Fig. 1A & 1B**). This indicates that Arr interacts with both ligands. To further confirm and quantitate the interactions of Arr with Rifampicin (positive control), c-di-AMP, and NAD+, we used microscale thermophoresis (MST). The MST titrations showed that Arr interacts with Rifampicin, NAD+, and c-di-AMP with the apparent dissociation constants (K_d_) of ∼6 nM, 100 nM, and 140 nM respectively (**Fig. 1C, 1D & 1E**). Together, NMR and MST titration experiments unambiguously showed that c-di-AMP interacts with Arr with a comparable affinity of NAD+ (co-factor), thus resulting in competitive inhibition of the enzyme and preventing ribosylation of rifampicin.

**Table 1.** List of MIC values (μg/ml.) of different strains against rifampicin.

| Strains | MIC (µg/ml) |
| --- | --- |
|  | Rifampicin |
| *M. Smegmatis* WT | 16 |
| *M. Smegmatis* Δ*pde* | 2 |
| *M. Smegmatis* Δ*pde*+pP*de* | 16 |
| *M. Smegmatis* Δ*disA* + pMV261 empty | 16 |
| *M. Smegmatis* Δ*disA* + pMV261-*disA* | 4 |
| *M. Smegmatis* Δ*disA* + pMV261-*disA* (D84A) | 16 |
| *M. Smegmatis* WT + pMV261-*arr* | 32 |
| *M. Smegmatis* Δ*pde* + pMV261-*arr* | 32 |
| *M. Smegmatis* Δ*arr* + pMV261 empty | 0.25 |
| *M. Smegmatis* Δ*arr* + pMV261-*disA* | 0.25 |
| *M. Smegmatis* Δ*arr* + pMV261-*disA* (D84A) | 0.25 |

**Figure 1.**
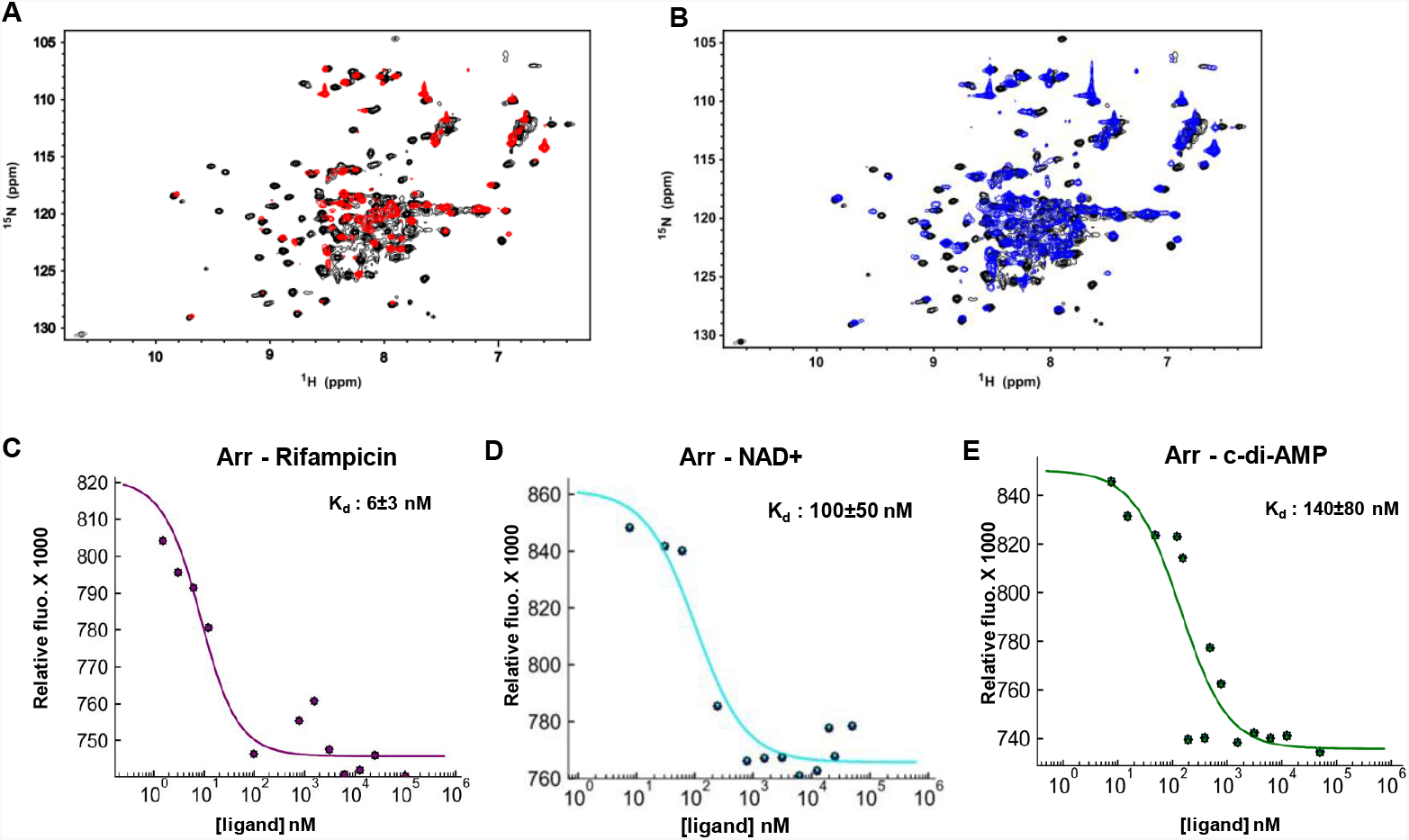
Biophysical enumeration of the interaction of Arr with c-di-AMP. NMR spectroscopic validation of interaction of Arr protein (A) with NAD+, where the overlay of 2D 1 H-15 N HSQC spectra of free Arr (black) and Arr with NAD+ (red) at 1:2 molar ratio is shown and (B) with c-di-AMP; an overlay of 2D 1 H-15 N HSQC spectra of free Arr (black) and Arr with c-di-AMP (blue) at 1:2 molar ratio is shown. Physical interaction of Arr with (C) rifampicin, (D) NAD+, and (E) c-di-AMP was confirmed by microscale thermophoresis (MST). Binding affinities in terms of dissociation constants (K_d_) are also mentioned in individual cases.

### LfrR protein expression is mediated by c-di-AMP resulting in ciprofloxacin sensitivity

To our surprise, we found that the Δ*disA* strain was moderately resistant to second-generation fluoroquinolone ciprofloxacin, which was proved in multiple assays. Our previous (disc inhibition assay) data (*10*) clearly showed that Δ*disA* strain had a small yet significant resistance phenotype against ciprofloxacin and with the help of mutants, we proved that the phenomenon is linked to c-di-AMP only, and not due to the absence of DisA enzyme’s DNA scanning (*10*) property. We checked the increased resistance level of Δ*disA* strain by minimum inhibitory concentration (MIC) assay, where the difference of ciprofloxacin concentrations from one well to the next well is 1.15 fold instead of the standard 2 fold so that we could precisely measure the MIC value. Our data showed a modest increase (∼2 fold) of MIC in Δ*disA* strain (0.869 μg/ml.) compared to WT (0.468 μg/ml.) (**Table 2**). Following this, we did an MBC (minimum bactericidal concentration) estimation of the strains and found the MBC value of Δ*disA* (1.75 μg/ml) strain was higher than WT (1 μg/ml) strain (**Table 2**). Similarly, when we compared the survival percentage of cells after treatment with 2.5 μg/ml. ciprofloxacin (10X MIC) for 24 hours, the killing profile of Δ*disA* strain looked different and showed a higher survival against ciprofloxacin compared to *M. smegmatis* WT (**Fig. 2A & 2B**). In all the cases, the complemented strain (Δ*disA*+pDisA) showed a reversal of resistance to the WT level and further confirmed the direct link of ciprofloxacin tolerability with the presence or absence of c-di-AMP (**Table 2**). Next, the EtBr efflux assay (**Fig. S2**) suggested that Δ*disA* strain had increased efflux pump activity which correlates well with the moderately low level of resistance phenotype, unlike a target gene mutation (*11*), where large shifts in MIC values are typically observed. Based on this, we shortlisted some candidate genes from the literature and one candidate that has drawn our attention was the LfrR-LfrA efflux module (*23, 24*). Once identified, we were specifically interested to know if the repressor gene *lfrR* was negatively downregulated by c-di-AMP. Indeed, our RNA-seq data revealed a significant fold change of expression (Log2 fold change: -1.378) of the *lfrR* repressor gene (*MSMEG_6223*, a TetR/AcrR family transcriptional regulator) in Δ*disA* strain; later the qRTPCR data also confirmed a significant downregulation of *lfrR* gene (**Fig. 2C**). Next, we overexpressed the *lfrR* gene from the pMV261 vector under a moderately strong constitutive *hsp60* promoter (P_*hsp60*_) and more importantly without the native 5’-UTR which might be responsible for c-di-AMP interaction, in Δ*disA* strain and found that the MIC value of *M. smegmatis* Δ*disA*+pLfrR strain dropped by ∼ 2 fold to the WT level (0.395 μg/ml.). In identical conditions, the control strain *M. smegmatis* Δ*disA*+pEmpty retained the higher MIC (0.869 μg/ml) (**Table 2**). Parallelly, ciprofloxacin (2.5 μg/ml) disc diffusion assay showed a significant increase in drug sensitivity for *M. smegmatis* Δ*disA*+pLfrR strain compared to *M.smegmatis* Δ*disA*+pEmpty strain (**Fig. 2D**). Both these observations further confirmed that c-di-AMP driven transcriptional downregulation of efflux pump repressor *lfrR* in *M. smegmatis*, which in turn results in the increased efflux of ciprofloxacin.

**Table 2.** List of MIC and MBC values (μg/ml.) of different strains against ciprofloxacin.

| Ciprofloxacin |  |
| --- | --- |
| Strains | MIC (µg/ml) |
| *M. Smegmatis* WT | 0.468 |
| *M. Smegmatis* Δ*disA* | 0.869 |
| *M. Smegmatis* Δ*disA*+pDisA | 0.468 |
| *M. Smegmatis* WT + pMV261 empty | 0.468 |
| *M. Smegmatis* Δ*disA* + pMV261 empty | 0.869 |
| *M. Smegmatis* WT + pMV261-*lfrR* | 0.395 |
| *M. Smegmatis* Δ*disA* + pMV261-*lfrR* | 0.395 |
| Strain | MBC (µg/ml) |
| *M. Smegmatis* WT | 1 |
| *M. Smegmatis* Δ*disA* | 1.75 |
| *M. Smegmatis* Δ*disA*+pDisA | 1 |

**Figure 2.**
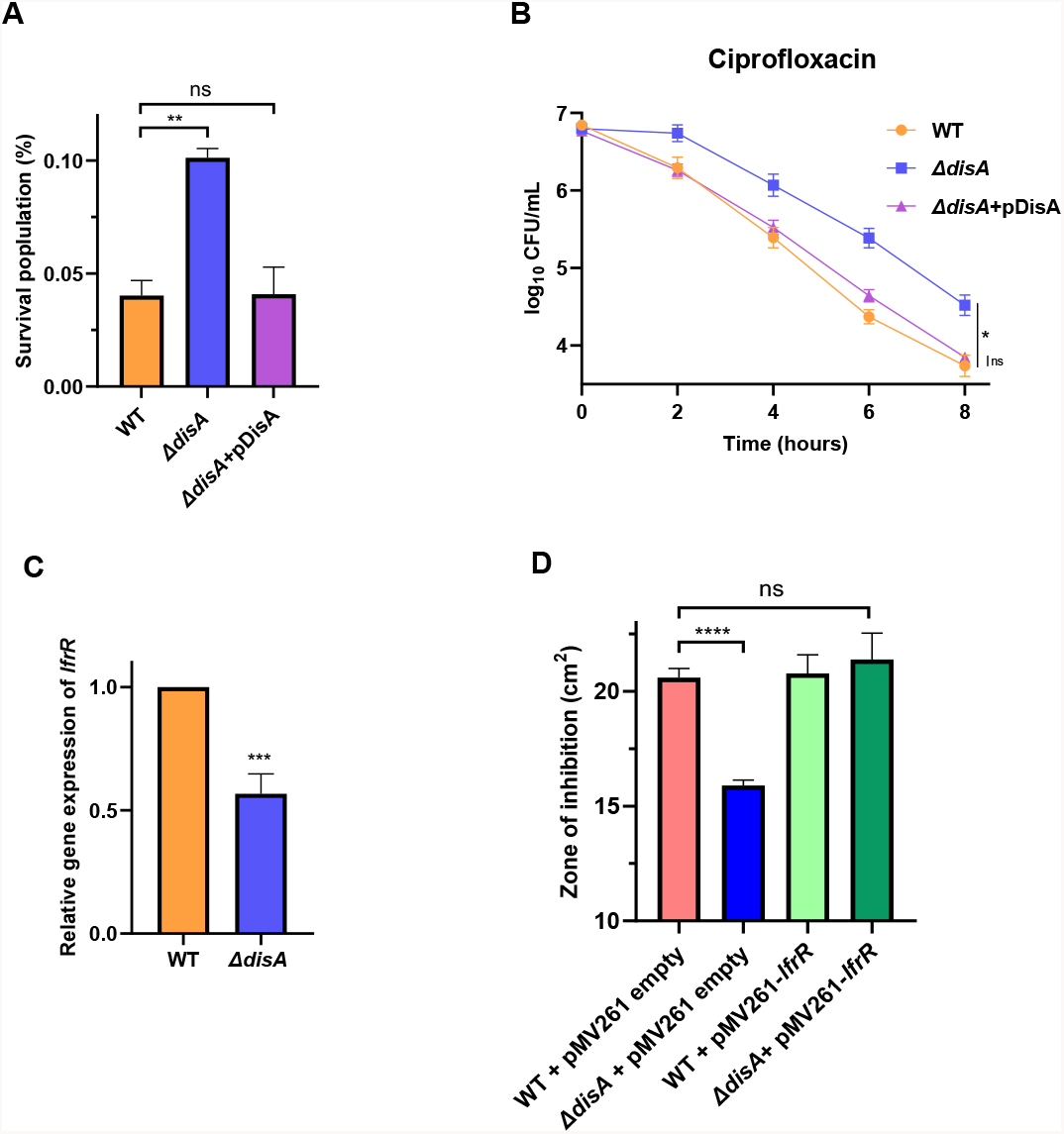
Elucidation of the role of c-di-AMP in ciprofloxacin resistance phenotype. The c-di-AMP null mutant strain *M. smegmatis* Δ*disA* shows more resistance to ciprofloxacin compared to *M. smegmatis* WT strain shown in multiple assays. (A) the survival percentage of *M. smegmatis* Δ*disA* strain was 2 fold higher than *M. smegmatis* WT strain after 2.5 μg/ml. ciprofloxacin (10X MIC) treatment for 24 hours, whereas the complemented strain Δ*disA*+pDisA showed a similar level of sensitivity to WT. (B) Time kill kinetics of *M. smegmatis* Δ*disA* strain shows moderate resistance to ciprofloxacin compared to WT and complemented strain. (C) qRT-PCR validation of significant downregulation of *lfrR* gene (repressor) in *M. smegmatis* Δ*disA* strain, confirming the basis of low-level ciprofloxacin resistance. (D) Ciprofloxacin disc diffusion assay shows that the overexpression of LfrR repressor from multicopy pMV261 vector reverses the ciprofloxacin resistance phenotype in *M. smegmatis* Δ*disA* strain. All the graphs are plotted using GraphPad Prism8, unpaired t-test was used to calculate statistical significance: ^***^ = P < 0.001; ^**^ = P < 0.01; ^*^ = P < 0.05; ns= non-significant.

### Several ribosomal proteins are downregulated in absence of c-di-AMP leading to increased susceptibility to different translation inhibitors

Our comprehensive analysis of basic phenotypes did not reveal any particular difference between *M.smegmatis* WT and Δ*disA* strain, except for the growth profile in minimal media (M9) and particularly in the presence of several protein synthesis inhibitors. In particular, we observed significant growth inhibition in the early exponential phase of growth of Δ*disA* strain in the presence of protein synthesis inhibitors erythromycin (**Fig. 3A**) and tobramycin (**Fig. 3B**) at 10 μg/ml and 0.2 μg/ml respectively. Next, we performed a MIC assay with erythromycin and found a ∼1.75 fold difference in MIC value between WT and Δ*disA* strain further confirming sensitivity phenotype against macrolide antibiotic erythromycin targeting 50S ribosomal subunit (**Table 3**). The complemented strain (Δ*disA*+pDisA) showed a reversal of sensitivity to the WT level and thus validated the bonafide connection of c-di-AMP with differential drug sensitivity. Our third assay which confirmed increased susceptivity of Δ*disA* strain was Combi-MIC when we measured the MIC of one drug in the presence of another drug by using the checkerboard method (*25*) for the following combinations: Erythromycin + Chloramphenicol (combination I) and Erythromycin + Streptomycin (combination II). To evaluate whether a particular combination resulted in an additive or synergistic outcome, we calculated the fractional inhibitory concentration index (ΣFIC) and found synergistic effects of the drug combinations in both strains. However, the significant difference in ΣFIC values (**Table 4**) indicated the increased susceptibility phenotype of Δ*disA* strain against two combined protein synthesis inhibitors. In a recent study (*10*), the differential expression of several ribosomal genes was highlighted by RNA-seq based transcriptome analysis. The three most significant downregulated ribosomal genes were found to be *rplL* (large protein L7/L12), *rpsR2* (small protein S18) and *rpsP* (small protein S16). Subsequently, we performed qRTPCR and confirmed significant downregulation *rplL* (**Fig. 3C**), which was earlier detected by RNA-seq analysis. To understand the biological effect possibly related to a defect in ribosomal biogenesis and/or assembly of subunits we performed a growth curve analysis in minimal media at a low temperature (20 °C) and it showed a clear growth defect in the Δ*disA* strain. Subsequently, we overexpressed *rplL* gene from multicopy pMV261 vector with constitutive hsp60 promoter and found that the low-temperature growth deficiency in Δ*disA* strain is recovered suggesting that the loss of L7/L12 protein was responsible for the growth defect in the Δ*disA* strain (**Fig. 3D**). However, no difference in ribosomal subunit concentration could be found when polysome profiling was performed from the WT and Δ*disA* strains, suggesting Δ*disA* strain probably did not have significant ribosome biogenesis defects (**Fig. S3A and S3B**).

**Table 3.** List of MIC values (μg/ml.) of different strains against erythromycin.

| MIC (µg/ml) |  |
| --- | --- |
| Strains | Erythromycin |
| *M. Smegmatis* WT | 48.4 |
| *M. Smegmatis* Δ*disA* | 27.7 |
| *M. Smegmatis* Δ*disA*+pDisA | 48.4 |

**Table 4.** A comparison of Combination MIC values of erythromycin-chloramphenicol and erythromycin-streptomycin combinations against *M. smegmatis* WT and Δ*disA* strains.

|  |  | MIC (µg/ml) |  | FIC (µg/ml) |  |
| --- | --- | --- | --- | --- | --- |
| Strain | Antibiotic | Alone | In combination | FIC | ΣFIC |
| *M. Smegmatis*<br>WT | Erythromycin | 32 | 2 | 0.062 | 0.125 |
|  | Chloramphenicol | 16 | 1 | 0.062 |  |
| *M. Smegmatis*<br>*ΔdisA* | Erythromycin | 32 | 1 | 0.031 | 0.046 |
|  | Chloramphenicol | 16 | 0.25 | 0.015 |  |
| *M. Smegmatis*<br>WT | Erythromycin | 32 | 8 | 0.25 | 0.75 |
|  | Streptomycin | 0.5 | 0.25 | 0.5 |  |
| *M. Smegmatis*<br>*ΔdisA* | Erythromycin | 32 | 4 | 0.125 | 0.375 |
|  | Streptomycin | 0.5 | 0.125 | 0.25 |  |

**Figure 3.**
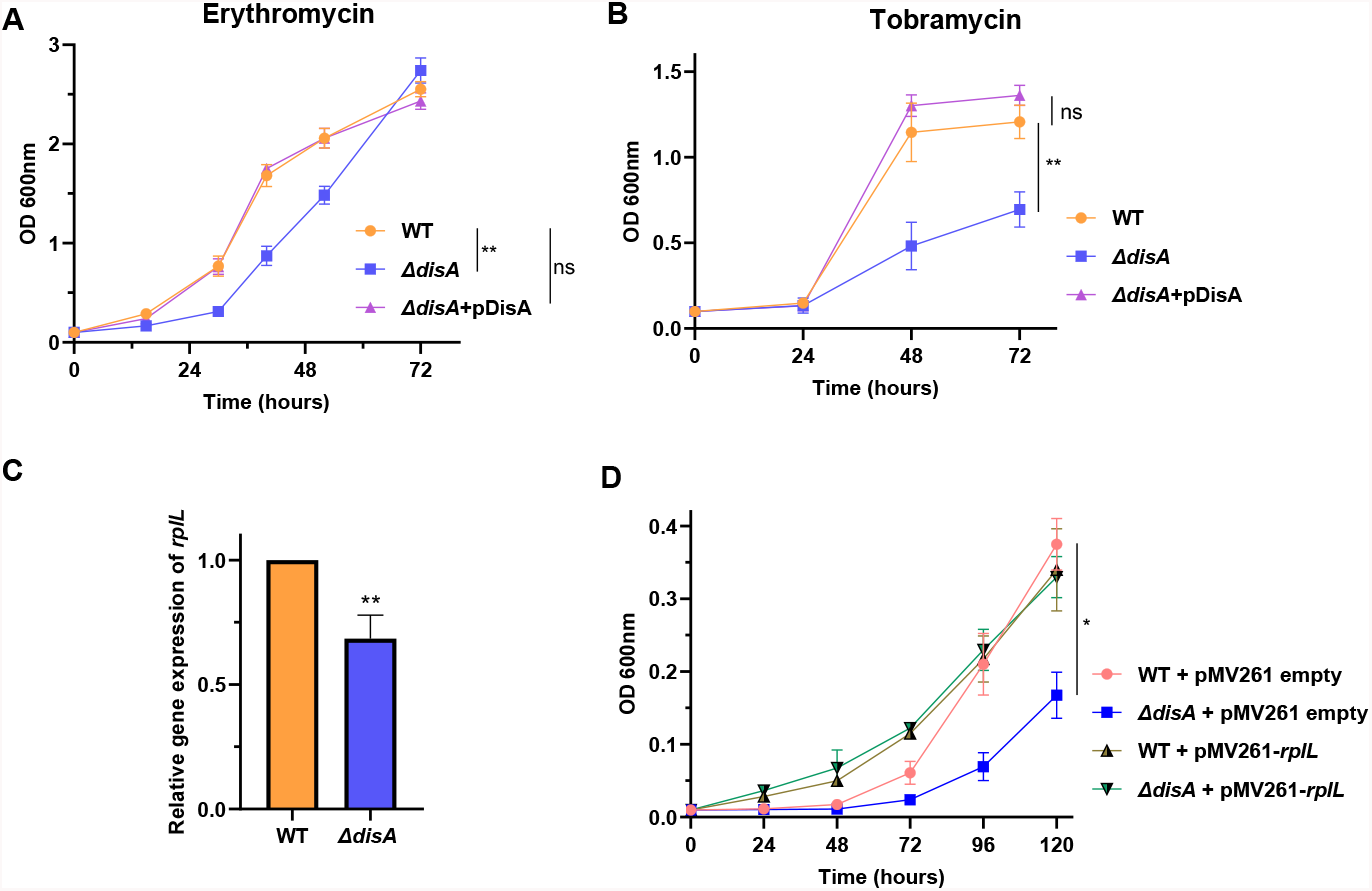
Lack of c-di-AMP makes Δ*disA* strain susceptible to multiple stresses. *M. smegmatis* Δ*disA* strain shown increased susceptibility against two different protein synthesis inhibitors (A) erythromycin (macrolide) and (B) tobramycin (aminoglycoside) pointing towards a possible defect in ribosome structure and/or activity in absence of c-di-AMP. (C) The qRT-PCR validation of significant downregulation of *rplL* gene (encodes for ribosomal protein L7/L12) in *M. smegmatis* Δ*disA* strain, reconfirming the observation from RNA-Seq analysis. (D) *M. smegmatis* Δ*disA* strain was shown to have a possible ribosome-related growth defect at low temperature (20°C), which was recovered by overexpression of RplL protein from multicopy pMV261 vector. All the graphs are plotted using GraphPad Prism8, unpaired t-test was used to calculate statistical significance: *** = P < 0.001; ^**^ = P < 0.01; ^*^ = P < 0.05; ns= non-significant.

### Δ*disA* ribosome shows compromised activity in protein synthesis

The L7/L12 protein is important for efficient and accurate translation (*26-29*). Since Δ*disA* ribosomes lead to under-expression of the L7/L12 proteins defect in translation could be anticipated. To identify the defects of the Δ*disA* ribosomes we tested those in a full-length protein (GFP+) synthesis assay (**Fig. 4A**) using a *M. smegmatis*-based fully reconstituted transcription-translation system (*30,31*). As a control, we used ribosomes from the genetic equivalent WT and *disA* complemented (Δ*disA*+pDisA) strains. Our results show that Δ*disA* ribosome is active in protein synthesis, but its activity is highly compromised as it could produce only 60% GFP+ protein compared to the WT in a given time of 1 hour (**Fig. 4B**). The ribosomes from (Δ*disA*+pDisA) strain largely rescued the defect as it increased GFP+ production to ∼85% (**Fig. 4B**).

**Figure 4.**
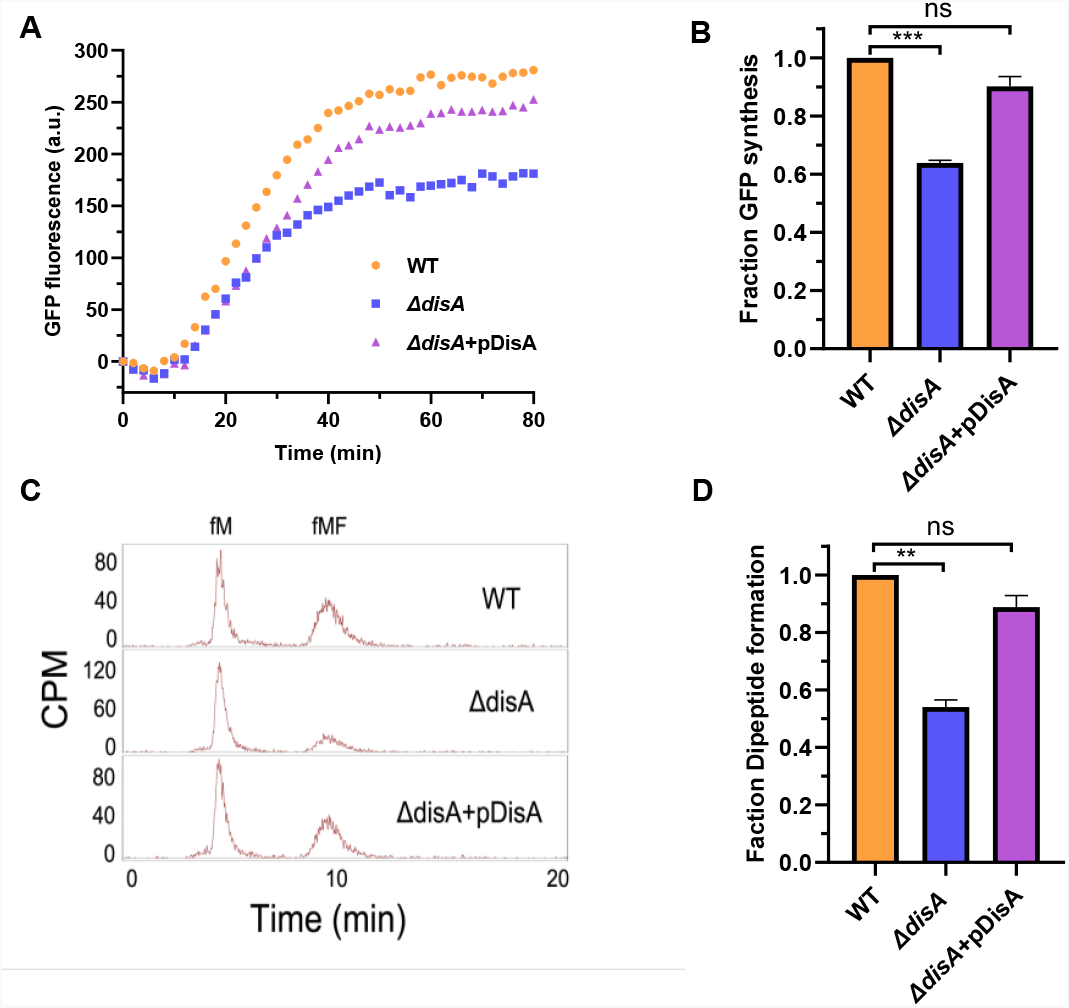
Compromised activity of the Δ*disA* ribosomes in protein synthesis. (A) Time course of GFP+ synthesis with Δ*disA*, WT and Δ*disA*+pDisA ribosomes in a *M. smegmatis*-based fully reconstituted transcription-translation system, followed with fluorescence readout (512 nm) in real-time. (B) The fraction of GFP+ synthesized in 1 hour with various ribosomes are labelled. (C) HPLC separation profiles of the mono (^3^H-fMet), and dipeptide (^3^H-fMet-Phe) with Δ*disA*, WT and Δ*disA* + pDisA ribosomes. (D) The fraction of dipeptide formed with the ribosome variants are indicated. All the graphs are plotted using GraphPad Prism8, unpaired t-test was used to calculate statistical significance: ^***^ = P < 0.001; ^**^ = P < 0.01; ^*^ = P < 0.05; ns= non-significant.

To check further whether the defect in Δ*disA* ribosomes originates from the main catalytic step of protein synthesis, we have tested those in a dipeptide synthesis assay, which includes both decoding and peptide bond formation steps. The relative amount of the [^3^H]-fMet monopeptide and [^3^H]-fMet-Phe dipeptides separated in reverse phase HPLC clearly shows less amount of (fMF) dipeptide formed with Δ*disA* ribosomes compared to the WT and Δ*disA*+pDisA ribosomes (**Fig. 4C**). Quantification of the dipeptide peaks suggests only ∼54% dipeptide formed with Δ*disA* ribosomes (**Fig. 4D**), which matches well with its capacity in total protein synthesis. The ribosomes from Δ*disA*+pDisA strain increased dipeptide production to ∼87%, also in good agreement with its activity in GFP+ synthesis (**Fig. 4D**). These results suggest that Δ*disA* ribosomes are primarily defective in the elongation step of protein synthesis, which leads to the defect in total protein synthesis.

## Discussion

We show here the physiological relevance of c-di-AMP in modulating the antibiotic resistance and sensitivity phenotypes of *M. smegmatis*. Our results of c-di-AMP messenger controlling drug resistance phenotypes of a single organism against four different classes of antibiotics is a unique observation. In addition to that, we have done a deep investigation to find out the underlying mechanisms behind the differential drug sensitivity in a case-by-case manner (**Fig. 5**). The fact that c-di-AMP interacts with Msm_Arr_ enzyme and thus inhibits its rifampicin ribosylation activity, targeting c-di-AMP degradation enzyme Pde (by which the cells will have an elevated intracellular c-di-AMP concentration) can be a logical approach to make the cells sensitive to rifampicin. Arr has been targeted before to augment rifampicin activity using specific libraries and an antibiotic-adjuvant combination has been shown to inhibit Arr that makes bacterial cells sensitive to rifampicin (*22*). Given this, our finding would be clinically significant to target ADP-ribosyltransferase MAB_0591 in pathogenic non-tuberculous bacteria like *M. abscessus* to make the organism more sensitive to rifampicin (*32*).

**Figure 5.**
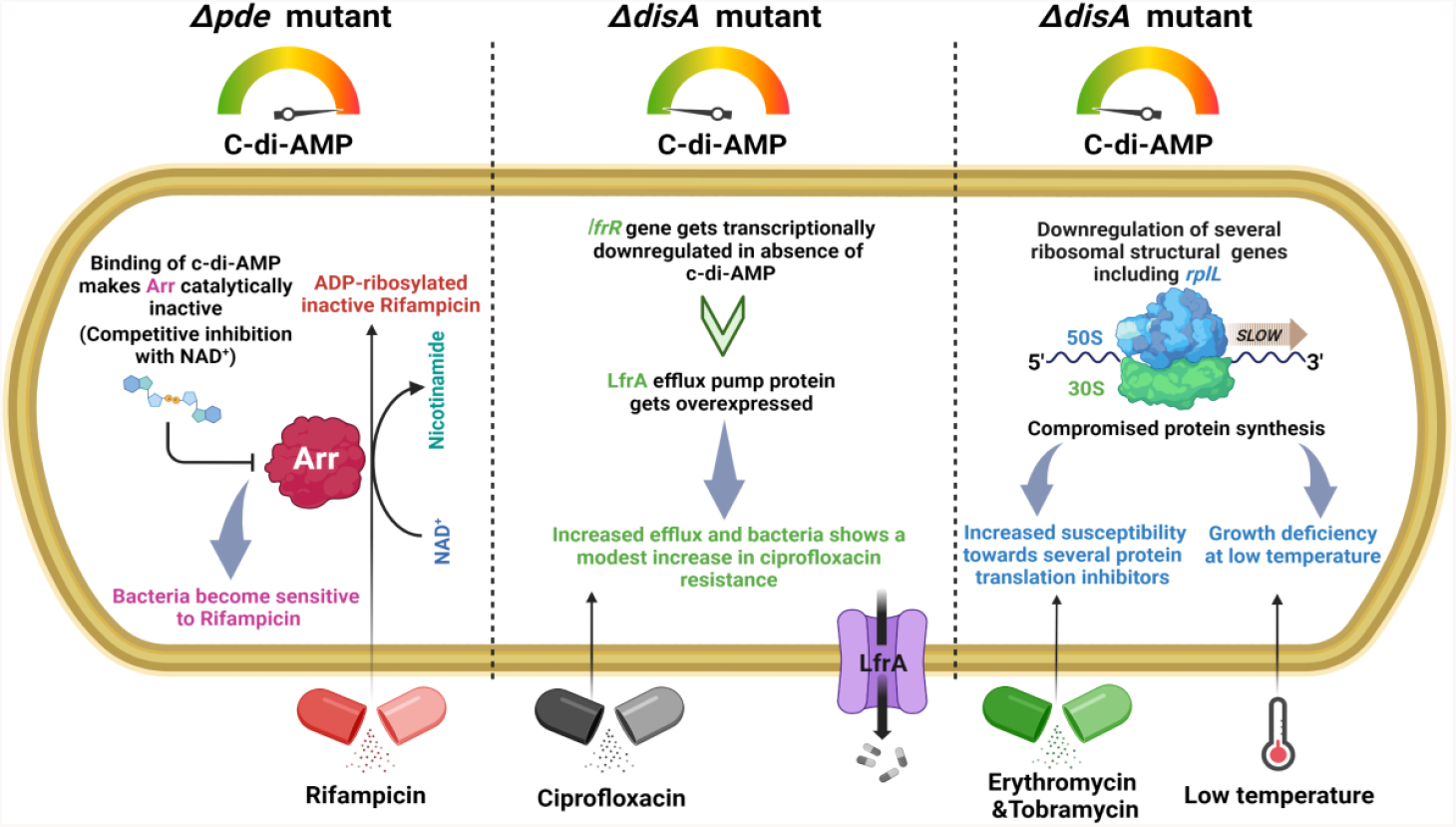
Schematic representation of our proposed model showing how c-di-AMP modulates multiple drug sensitivity pathways in *M. smegmatis* (created with BioRender.com).

One important finding of our study is the intrinsic modulation of efflux pump activity linked to intracellular c-di-AMP concentration in *M. smegmatis*, which could be a survival strategy under transient fluoroquinolone exposure without accumulating any target mutation, especially when such mutation often comes with a huge fitness cost *(11)*. Another second messenger ppGpp was also shown recently to modulate efflux pump-related gene expression in *A. baumannii* (*33*) and result in MIC modulation of several antibiotics. Further, we checked the presence of possible riboswitch or secondary structure upstream of the *lfrR* gene by bioinformatics analysis. We identified a putative stem-loop structure at the -35 region upstream of the *lfrR* gene (data not shown), which possibly blocks RNA polymerase’s access to the promoter necessary to drive transcription of LfrR. We might need to study this in detail in the future.

Finally, we reported that the lack of c-di-AMP significantly downregulates the expression of several ribosomal proteins in *M. smegmatis*, which leads to a compromised activity of the ribosomes in protein synthesis, hypersensitivity of the strain to low temperature and to several antibiotics targeting protein synthesis (translation) machinery. The Δ*disA* ribosomes are defective in the elongation step of protein synthesis, which makes the total protein synthesis relatively slower. Probably the slower protein synthesis widens up the time window for binding of the antibiotics to the Δ*disA* ribosomes thereby making those more sensitive to the ribosome targeting antibiotics. A second messenger molecule modulating ribosome structure and function is a novel observation in any bacteria and would possibly open up a new field or research in the context of targeting such auxiliary pathways such as c-di-AMP synthesis for novel antibiotic research.

Thus, our study has delineated the specific genes and pathways regulated by c-di-AMP in the transcriptional and post-translational levels. Though c-di-AMP was discovered many years back (*34*), an elaborated study of this novel second messenger’s important role in dictating antibiotic resistance and sensitivity phenotypes in a mechanism-driven manner was still lacking. The results presented in this paper related to c-di-AMP driven differential antibiotic sensitivity response mechanisms in the model organism *M. smegmatis* gave novel insights which will provide further advancement in this field of second messenger signaling research. Since the c-di-AMP has been shown to have a definite link with *M. tuberculosis* virulence and triggering an immune response of the host (*35, 36*), this study on the model organism *M. smegmatis* could give an ideal starting point to shape the future research related to the emergence of drug-resistant strains of *M. tuberculosis*. A detailed study in the future should be aimed to probe c-di-AMP’s role in the modulation of antibiotic sensitivity and resister development in *M. tuberculosis*, which might help understand the dynamics of resistant mutant enrichment and subsequently design a superior treatment strategy to treat MDR *M. tuberculosis*.

## Acknowledgment

We thank Prof. Dipankar Chatterji and Prof. Umesh Varshney at Indian Institute of Science, Bangalore for valuable feedback on the work. Authors acknowledge the DST and DBT, India, for the NMR and MST facilities at the Indian Institute of Science, Bangalore.

## Author contributions

A.P. and A.G. conceived the study. A.P., S.S., M.S. and A.G. designed the research; A.P., D.G., B.S., X.G., A.S., V.C. and A.G. performed experiments; A.P., S.S., U.V., M.S. and A.G. analysed the data; A.P., S.S., M.S. and A.G. wrote the paper.

## Data and materials availability

All data are available in the manuscript or the supplementary materials.

## List of Supplementary Materials

Materials and Methods.

Figs. S1 to S3.

Table 1 (bacterial strains) and Table 2 (primers).

References (37-49): specific to materials and methods which are not present in the main text.

