## Supplementary data for "Second messenger c-di-AMP regulates multiple antibiotic sensitivity pathways in *Mycobacterium smegmatis* by discrete mechanisms"

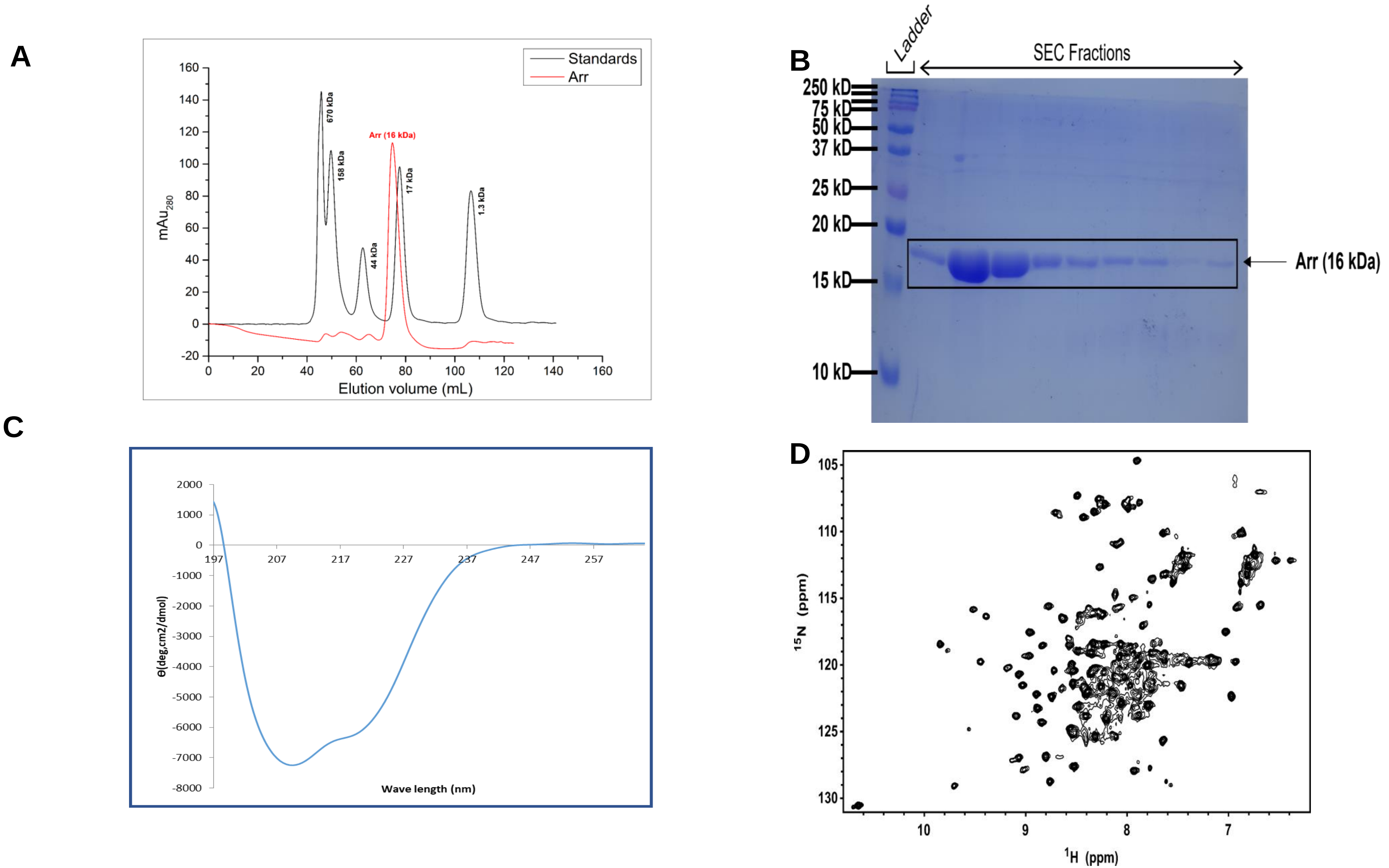


**Figure S1:** (A) SEC elution profile of Arr protein, overlayed with the elution profile of standard proteins, showing that Arr eluted as a monomer; (B) SDS-PAGE analysis of Arr elution fractions from SEC, indicating the protein is purified to near homogeneity ; (C) CD spectrum of Arr indicating that Arr is α-helical in solution. (D) 2D 1 H-15 N HSQC NMR spectrum of free Arr with good chemical shift dispersion suggesting that protein is folded in solution.

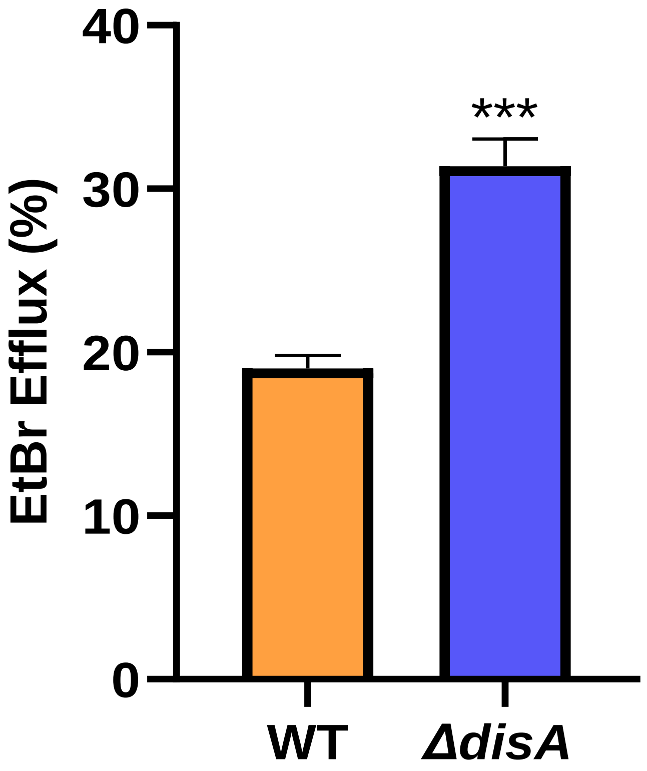


**Figure S2:** *M. smegmatis* Δ*disA* strain shows increased Ethidium bromide (EtBr) efflux activity.

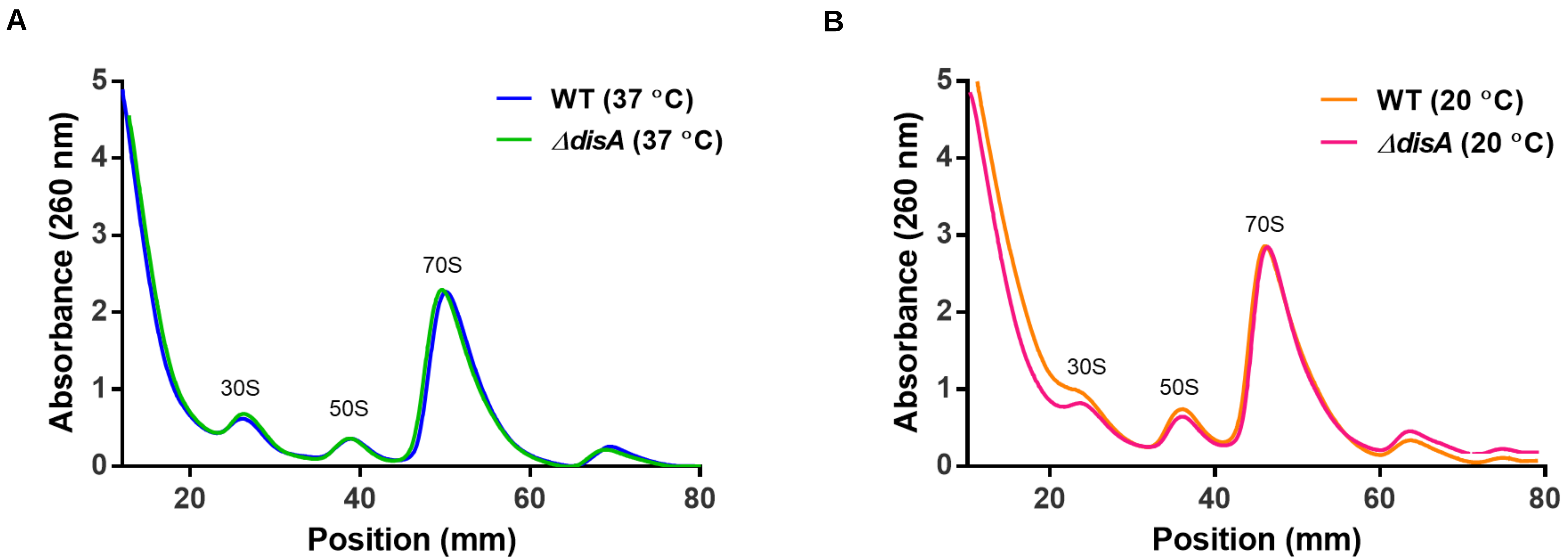


**Figure S3:** Polysome profiling reveals no conclusive difference in ribosomal subunits' profile of (A) Normal (37°C) and (B) low (20°C) temperature grown cultures of *M. smegmatis* WT and Δ*disA* strain.

**Table S1:** Bacterial strains used in the study

| Strains | Description | Source/ Reference |
|---|---|---|
| *M. smegmatis* Wild type (WT) | *M. smegmatis* mc$^2$ 155 | Laboratory stock |
| *M. smegmatis* Δ*disA* | *M. smegmatis* mc$^2$ 155 strain with knock out of *disA* ; Kan$^R$ | Ref. 10 |
| *M. smegmatis* Δ*pde* | *M. smegmatis* mc$^2$ 155 strain with knock out of *pde ;* Kan$^R$ | Ref. 10 |
| *M. smegmatis* Δ*disA* +*p*DisA | *M. smegmatis* Δ*disA* strain with *disA* cloned in pMV361 vector; Hyg$^R$,Kan$^R$ | Ref. 10 |
| *M. smegmatis* Δ*pde* +*p*Pde | *M. smegmatis* Δ*pde* strain with *pde* cloned in pMV361 vector; Hyg$^R$,Kan$^R$ | Ref. 10 |
| *M. smegmatis* Δ*disA* +pMV261-*disA* (D84A) | *M. smegmatis* Δ*disA* strain with *disA* (D84A mutant) cloned in pMV261 vector; Hyg$^R$,Kan$^R$ | This study |
| *M. smegmatis* Δ*disA* +pMV261-*disA* | *M. smegmatis* Δ*disA* strain with *disA* cloned in pMV261 vector; Hyg$^R$,Kan$^R$ | This study |
| *M. smegmatis* WT + pMV261 empty | *M. smegmatis* WT strain with pMV261 empty vector; Hyg$^R$ | Ref. 11 |
| *M. Smegmatis* Δ*disA* +pMV261 empty | *M. smegmatis* Δ*disA* strain with pMV261 empty vector; Hyg$^R$ ,Kan$^R$ | This study |
| *M. smegmatis* Δ*pde* +pMV261 empty | *M. smegmatis* Δ*pde* strain with pMV261 empty vector; Hyg$^R$,Kan$^R$ | Ref. 11 |
| *M. smegmatis* Δ*arr* +pMV261 empty | *M. smegmatis* Δ*pde* strain with pMV261 empty vector; Hyg$^R$,Kan$^R$ | This study |

**Table S1:** Bacterial strains used in the study (cont.)

| **Strains** | **Description** | **Source/ Reference** |
|---|---|---|
| *M. smegmatis* WT + pMV261-*lfrR* | *M. smegmatis* WT strain with *lfrR* cloned in pMV261 vector; $Hyg^R$ | This study |
| *M. smegmatis* Δ*disA* + pMV261-*lfrR* | *M. smegmatis* Δ*disA* strain with *lfrR* cloned in pMV261 vector; $Hyg^R$,$Kan^R$ | This study |
| *M. smegmatis* WT + pMV261-*rplL* | *M. smegmatis* WT strain with *rplL* cloned in pMV261 vector; $Hyg^R$ | This study |
| *M. smegmatis* Δ*disA* + pMV261-*rplL* | *M. smegmatis* Δ*disA* strain with *rplL* cloned in pMV261 vector; $Hyg^R$,$Kan^R$ | This study |
| *M. smegmatis* WT + pMV261-*arr* | *M. smegmatis* WT strain with *arr* cloned in pMV261 vector; $Hyg^R$ | This study |
| *M. smegmatis* Δ*pde* + pMV261-*arr* | *M. smegmatis* Δ*pde* strain with *arr* cloned in pMV261 vector; $Hyg^R$,$Kan^R$ | This study |
| *M. Smegmatis Δarr* + pMV261-*disA* | *M. smegmatis Δarr* strain with *disA* cloned in pMV261 vector; $Hyg^R$,$Kan^R$ | This study |
| *M. Smegmatis Δarr* + pMV261-*disA* (D84A) | *M. smegmatis Δarr* strain with *disA* (D84A mutant) cloned in pMV261 vector; $Hyg^R$,$Kan^R$ | This study |
| *E. coli* BL21 + pET28a-*arr* | *E. coli* BL21 Rosetta (DE3) cells with *arr* overexpressed in pET28 vector; $Kan^R$ | This study |

**Table S2:** Primers used in the study

| **Name** | **Sequence (5' to 3')** | **Description** |
|---|---|---|
| pET28_arr_REV_NotI | ACTGGCGGCCGCGTCATAGATGACCGCCAG | To construct pET28a-*arr* |
| pET28_arr_FW_NcoI | CATGCCATGGGGGCGAATCCGCCGAAACCG | To construct pET28a-*arr* |
| lfrR_FW_BamHI | ctagGGATCCatgaccagcccgagcatcga | To construct pMV261-*lfrR* |
| lfrR_REV_HindIII | actgAAGCTTtcaggtgcgcggcagggtga | To construct pMV261-*lfrR* |
| fw_pmv261H_rplL | AGCTGGATCCtcaggaaggaccataaacat | To construct pMV261-*rplL* |
| rev_pmv261H_rplL | TCAGAAGCTTactacttgacggtgaccgag | To construct pMV261-*rplL* |
| sigA mid FW | cgccgagaagggcgagaagc | For performing RT-PCR of *sigA* |
| sigA mid REV | tcaggcccaggttgccttcc | For performing RT-PCR of *sigA* |
| lfrR new mid fw | catcgtcgaggtgctcaac | For performing RT-PCR of *lfrR* |
| lfrR new mid rev | gtccttggctgcctcataac | For performing RT-PCR of *lfrR* |
| rplL mid fw | cggtcgcggttgccgctgcc | For performing RT-PCR of *rplL* |
| rplL mid rev | gggcgcgctgtcgacgagat | For performing RT-PCR of *rplL* |
