## Supplementary information for "Second messenger c-di-AMP regulates multiple antibiotic sensitivity pathways in *Mycobacterium smegmatis* by discrete mechanisms"

### Supplementary material

#### Materials and Methods:

##### ***Bacterial strains, media and growth conditions***

*M. smegmatis* mc$^2$155 (WT), it's knockout variants *ΔdisA & Δpde* and the overexpression constructs were grown in Middlebrook 7H9 broth (MB7H9; HiMedia) with 2% (wt/vol) glucose as a carbon source and 0.05% (vol/vol) tween-80, Agar (1.5%, w/v) (HiMedia) at 37°C. The antibiotics kanamycin and hygromycin were used at a concentration of 25 μg/ml and 50 μg/ml respectively (*10*). Ciprofloxacin, erythromycin and tobramycin powder were obtained from Sigma-Aldrich, USA and Sisco Research Laboratories.

For the growth inhibition assay, bacteria were grown in MB7H9 till the mid-exponential phase, washed twice and inoculated into M9 minimal media containing 1mM $MgCl_2$, 0.3mM $Cacl_2$ with 0.2% glucose (*37*) by adjusting the final optical density at 600 nm ($OD_{600}$) of 0.03. 5 ml. of such cultures were taken in a sterile glass tube. Either erythromycin or tobramycin was added to the specific culture tubes at a final concentration of 10 μg/ml and 0.2 μg/ml respectively and kept at 37°C shaking for 72 hours. The bacterial growth was further monitored in equal intervals by recording $OD_{600}$ and plotted using Graph Pad Prism Software. For the low-temperature growth assay, bacterial cultures were kept at 20°C shaking for 120 hours $OD_{600}$ was measured in equal intervals. All the growth assays were performed in a set of 3 replicates.

##### ***Minimum Inhibition Concentration (REMA) assay***

Minimum Inhibition Concentration (MIC) values of the antibiotics were estimated using REMA (Resazurin microtiter assay) adapted from an earlier protocol (*38*). In brief, the transmittance of the cultures was adjusted to a McFarland turbidity standard of 1 and then diluted to 1:10. Next, 196 µl of the diluted culture of respective strains were inoculated into 96 well microtiter plate wells containing a 2-fold or sometimes 1.15-fold (if we needed to find minor differences in MIC), serial dilution of the antibiotics of a specific range of concentrations. Plates were sealed and incubated at 37$^{o}$C for 36 hours. Next, 30 µl of sterile 0.01% resazurin dye was added to each well, and the plates were further incubated for 5-6 hours. The color of the resazurin changed from blue to pink in case of bacterial growth in wells. The MIC was determined as the minimum antibiotic concentration at which the resazurin did not change its color.

***Minimum bactericidal concentration***

To determine the minimum bactericidal concentration (MBC) of the drugs against different strains, bacteria were grown in MB7H9 media till the exponential phase and adjusted the final $OD_{600}$ to 0.2 in fresh MB7H9 medium. 4 ml. of such culture was added to four different tubes with variable concentrations of ciprofloxacin i.e. 0.5, 1, 1.5 and 2 µg/ml. (final concentration). Tubes were incubated at 37°C shaker and CFU estimations were done after 24 hours. Plates were further incubated at 37°C for 3-4 days for colony growth and subsequently counted. The minimum drug concentration which was able to kill 99.9% of the bacterial population (3 logarithmic decrease) within 24 hours was recorded as the MBC of ciprofloxacin against that particular strain.

***in vitro combination studies in broth***

The combination MIC involving two antibiotics was performed using the checkerboard method as mentioned by Balasubramanian et al. (*39*) Briefly, using 96-well microtiter plates, we added 2 µl of the first drug vertically and 2 µl of the second drug horizontally (from the antibiotic strength of 100 times) to obtain multiple combinations of these two drugs. Bacterial cultures were adjusted to

McFarland turbidity standard of 0.1 and 196 µl of such cultures were added to each well. The plates were incubated at 37°C for 36 hours. After that, 30 µl of 0.01% resazurin dye was added to each well, and the plates were further incubated for 5-6 hours. To evaluate whether the combinations of drugs have some additive or synergistic effect, we calculated the fractional inhibitory concentration index (ΣFIC) using this formula: ($MIC_{drug\ A}$ in combination/ $MIC_{drug\ A}$ alone) + ($MIC_{drug\ B}$ in combination/ $MIC_{drug\ B}$ alone).

##### *Disc diffusion assay*

As previously described (*40*), cultures were grown until the mid-exponential phase and adjusted the final $OD_{600}$ to 0.7 in fresh MB7H9 medium. 100µl of the culture was spread on MB agar plates and dried. Then, ciprofloxacin (2.5 µg/ml) discs were kept in the center of the plate. Plates were kept for incubation at 37°C for 3 days and the zone of inhibition ($cm^2$) was measured to calculate the antibiotic sensitivity.

##### *Time-kill kinetics assay*

*M. smegmatis* cultures were grown till the late-exponential phase ($OD_{600}$ of 1.3-1.5), diluted 1:100 to start a secondary culture and grown up to the mid-exponential phase ($OD_{600}$ of 0.6-0.8), and adjusted the final $OD_{600}$ to 0.2 corresponding to ~2 $X10^7$ CFU/ml in fresh MB7H9 medium. 4 ml of such culture was added to a glass tube and 2.5 µg/ml. (final concentration) ciprofloxacin (10X MIC) was added and CFU/ml was estimated by spreading it into the MB7H9 agar plate. Tubes were incubated at 37°C shaker and CFU estimations (plating) were done every 2 hours up to 8 hours. Plates were further incubated at 37°C for 3 days. This assay was performed with 3 biological replicates.

##### ***EtBr efflux Assay***

The efflux assay protocol was adapted from the previously described method (*11*,*41*). Briefly, *M. smegmatis* cultures were grown at 37°C in MB7H9 medium to $OD_{600}$ of 0.8-1. Cells were then washed twice with phosphate buffer saline (PBS) with 0.05% tween80, resuspended in 1/3$^{rd}$ volume of the PBS kept at 37°C shaking for one hour of starvation. After that, CCCP (carbonyl cyanide m-chlorophenylhydrazone, 100 μM final concentration) was added and the cells were further incubated for 30 minutes in the same condition. After that, ethidium bromide (0.5 μg/ml final concentration) was added and cells were incubated for another 30 minutes. Cells were then washed with the 1XPBST in the same volume to remove the CCCP and extracellular EtBr. After that, cells were taken into 96 well black plates and 2% glucose was added to the wells to enhance efflux activity. For quantifying the efflux of EtBr, fluorescence measurements were taken for the next one hour (with a 1 minute interval within two readings) in the Varioskan Flash multimode reader (Thermo Fisher Scientific) at 530 nm and 590 nm wavelengths for excitation and emission respectively.

##### ***Quantitative real-time PCR (qRT-PCR) experiment***

As previously described (*42*), secondary cultures of *M. smegmatis* were grown till the mid-exponential phase ($O.D._{600nm}$ 0.07-0.8) in the MB7H9 medium containing 2% glucose, and 0.05% tween. Cultures were then washed with 1X PBS and resuspended in Trizol (Sigma) and the cells were lysed by the bead-beating method. Qiagen's RNeasy Minikit was used for the RNA isolation. A one-step TB Green RT-PCR kit (Takara) was used to perform the qRT-PCR assay for the gene of interest and the assay was performed in Bio-Rad CFX Opus 96 Real-Time PCR system. Gene expression analysis was carried out using Bio-Rad CFX Maestro software and RNA levels were normalized to the *M. smegmatis sigA* (housekeeping) gene. The experiments were performed in three biological replicates and every time in technical duplicates.

***Expression and purification of Arr***

For Arr overexpression, the pET28-*arr* plasmid was transformed into *Escherichia coli* BL21 Rosetta (DE3) cells which were then cultured at 37˚C until the $OD_{600}$ reached 0.7. The cells were induced for protein expression with 1mM IPTG at 26˚C for 16 hrs. The cells were pelleted by centrifugation at 6,000 rpm for 15 mins. Sonication was used to lyse the cells resuspended in lysis buffer (20 mM Tris, 300 mM NaCl, pH 6.5). The lysate was centrifuged at 13,000 rpm for 90 mins to pellet down the cell debris. For Arr protein purification, the supernatant collected was loaded onto a 5 ml Ni2+-NTA Sepharose affinity column (GE Healthcare) at 4°C with a flow rate of 0.5 ml/min for binding. After the binding step, wash buffer (20 mM Tris, 300 mM NaCl, 20 mM imidazole, pH 6.5) was passed through the column at a 2 ml/min flow rate to remove non-specifically bound proteins. The bound Arr protein was eluted using elution buffer (20 mM Tris, 300 mM NaCl, 250 mM imidazole, pH 6.5) and passed at a 0.5 ml/min flow rate. The elution fractions were analyzed for the protein of interest using SDS–PAGE. Fractions containing the proteins were pooled and concentrated, and further loaded onto a Superdex 75 column (GE Healthcare) for size exclusion chromatography (SEC) purification using SEC buffer (20 mM Tris, 150 mM NaCl, pH 6.5). For overexpression of uniformly $^{15}N$ labeled Arr, cells were cultured in M9 minimal media containing one g/L of $^{15}NH4Cl$ till ~0.8 $O.D._{600nm}$, 0.5 mM Isopropyl β-d-1-thiogalactopyranoside (IPTG) was used for induction at 16°C for 16 hours. The final protein sample for NMR acquisition was prepared in 20 mM $NaH_2PO_4$, 200 mM NaCl, pH 6.5 buffer.

***Circular Dichroism (CD) spectroscopy***

CD spectroscopy was used to evaluate the secondary structure of free Arr in solution. The spectra were recorded on a JASCO J-715 spectropolarimeter using a 1 cm path length cuvette (Hellma Analytics) at 25°C at a scanning speed of 100 nm/min with a response time of 4s. The wavelength scan was done from 190 to 260 nm. A sample concentration of 15 μM was used for the experiment. The average of three iterations was taken for each sample, followed by baseline correction to negate the contribution from the buffer.

***Nuclear magnetic resonance (NMR) spectroscopy***

2D $^{1}$H-$^{15}$N HSQC NMR spectrum of Arr was recorded on Bruker 700 MHz, equipped with room temperature probe at 298 K. Arr sample was concentrated to 200 μM and 10% D20 was added for spectrometer deuterium lock. For the titration experiments, increasing amounts of unlabelled compounds (NAD+ and cyclic-di-AMP) were added to a $^{15}$N labelled Arr solution and a 2D $^{1}$H-$^{15}$N HSQC spectrum was recorded at each step of titration. Data were processed using Bruker TOPSPIN 3.1 and analyzed using NMRFAM-SPARKY (*43*).

***Microscale thermophoresis (MST)***

Binding affinity experiments were carried out on a Monolith NT.115 series instrument (Nano Temper Technologies GMBH). Arr protein was labeled with Monolith Protein labeling kit RED-NHS second-generation amine (Nano Temper Technologies GmbH) according to the manufacturer's guidelines. Roughly 10 μl of sample in MST buffer (20 mM HEPES [pH 6.5], 200 mM NaCl) were loaded into Monolith NT.115 regular capillaries, and thermophoresis was measured for 30 seconds. Analysis was performed with Monolith software. $K_d$ was quantified by analyzing the change in normalized fluorescence ($F_{\mathrm{norm}}$, fluorescence after thermophoresis/initial fluorescence) as a function of binding concentration. Curves for $K_d$ data were fitted to a 4-parameter logistic equation using nonlinear regression in Palmist 1.5.6 software (*44*) and plotted in Gussi software v1.1.0 (*45*).

***Polysome profiling***

The method for the polysome profiling was adapted from (*46*). Briefly, *M. smegmatis* culture was grown in M9 media at 37°C or 20°C till ~0.6 $O.D._{600nm}$ and chilled on the ice-salt mix. The culture was harvested at 7,000 rpm, 4°C for 5 min. Pellet was washed and resuspended in polysome buffer (20 mM Tris-HCl pH-8, 10 mM $MgCl_2$, 60 mM $NH_4Cl$, 3 mM DTT and 5% sucrose). The cell extract was prepared by beads beating and the lysate was centrifuged at 13,000 rpm, 4°C for 40 min. Approximately 10-15 $O.D._{260nm}$ of cell extract was loaded on a linear sucrose gradient (10-35%) prepared using BioComp gradient master. Samples were centrifuged using a Beckman SW41 rotor at 36,000 rpm, 4°C for 3.5 hours. Gradients were fractionated using a BioComp gradient fractionator with continuous $A_{260}$ measurement.

***in-vitro GFP protein synthesis assay***

70S ribosomes were purified from *M. smegmatis* WT, Δ*disA* and Δ*disA*+pDisA strains following the laboratory protocol (*47*). *M. smegmatis* initiation factors IF1, IF2, and IF3 and elongation factor EF-Tu and EF-G were cloned in pET24a plasmid with (His)x6-tag in the C-terminus, overexpressed in *E. coli* BL21-CodonPlus (DE3)-RIPL strain and purified to homogeneity using Ni-NTA affinity column connected to the ÄKTA-prime plus chromatography system *(30)*. For the *in vitro* protein synthesis assay, a reconstituted transcription-translation system was constructed that is composed of highly pure and active translation components from *M. smegmatis*, including 70S ribosomes, translation factors, bulk tRNA, and aminoacyl tRNA synthetases *(30)*. The reaction also contained amino acids, ATP, GTP, T7 RNA polymerase and an optimized energy-regenerating system containing phosphoenol pyruvate (PEP), pyruvate kinase and myokinase in HEPES-polymix buffer, pH 7.5 *(26)*. GFP synthesis was initiated by addition of pET24-GFP plasmid to the reaction mix at 37 ºC. Synthesis of active GFP protein was followed by monitoring GFP fluorescence (512 nm) with time in a TECAN Infinite 200 PRO plate reader (*47,48*).

***Dipeptide formation assay***

For MF dipeptide formation assay, the initiation complex and an elongation mix were prepared. The initiation complex was formed by mixing 2 μM 70S ribosome, 10 μM MFF mRNA (transcribed *in vitro* with T7 RNA polymerase and purified with Sephadex G-75 gel filtration chromatography), 1 μM $[^3H]$-fMet-$tRNA^{fMet}$ , charged and purified *(48)* and 2 μM of all three initiation factors in HEPES-polymix buffer, pH 7.5. An elongation mix was prepared by incubating 40 μM $tRNA^{Phe}$, 0.5 mM Phe, and 1 unit $^{Phe}$tRNA synthetase and 40 μM EF-Tu in the same buffer. The initiation complex and elongation mix were incubated separately at 37 ºC for 15 min. Dipeptide formation was initiated by mixing the two and the reaction was quenched after 10 sec by adding 17% formic acid. The $[^3H]$-fMet and $[^3H]$-fMet-Phe peptides were separated by reverse phase HPLC equipped with in line with radioactivity detection (β-RAM model 3 IN/US systems) as described by Wang et al. *(49)*.
